# A chemosensory switch couples genetic sex to behavioral valence

**DOI:** 10.1101/231936

**Authors:** Kelli A. Fagan, Jintao Luo, Ross C. Lagoy, Frank C. Schroeder, Dirk R. Albrecht, Douglas S. Portman

## Abstract

As a fundamental dimension of internal state, biological sex modulates neural circuits to generate naturally occurring behavioral variation. Understanding how and why circuits are tuned by sex can provide important insights into neural and behavioral plasticity. Here, we find that sexually dimorphic behavioral responses to *C. elegans* ascaroside sex pheromones are implemented by the functional modulation of shared chemosensory circuitry. In particular, the sexual state of a single sensory neuron pair, ADF, determines the nature of an animal's behavioral response regardless of the sex of the rest of the body. Genetic feminization of ADF causes males to be repelled by, rather than attracted to, ascarosides, while masculinization of ADF is sufficient to make ascarosides attractive to hermaphrodites. Genetic sex modulates ADF function by tuning chemosensation: ADF is able to detect the ascaroside ascr#3 only in males, a consequence of cell-autonomous action of the master sexual regulator *tra-1*. Genetic sex regulates behavior in part through the conserved DMRT gene *mab-3*, whose male-specific expression in ADF promotes ascaroside attraction. The sexual modulation of ADF has a key role in reproductive fitness, as feminization or ablation of ADF renders males unable to use ascarosides to locate mates. These results demonstrate that DMRT genes can functionally modulate shared neural circuits; moreover, they reveal an adaptive mechanism in which chromosomal sex controls a cell-autonomous switch that tunes sensory function, determines behavioral valence, and promotes reproductive fitness.

## INTRODUCTION

Sex differences in behavior provide a powerful framework for identifying mechanisms that sculpt naturally occurring behavioral variation [1-4]. These differences exist along a remarkably wide spectrum, ranging from innate, sex-specific, completely divergent behavioral programs, to subtle, population-level tendencies to behave differently under a given set of conditions [5]. Importantly, simpler model systems afford the advantage that the effects of biological sex on neural development and function can be largely (though see[6]) dissociated from the social influences that can cause apparent sex differences in more complex animals. In both *Drosophila* and *C. elegans*, biological sex regulates multiple aspects of neuronal development, including neurogenesis and neuronal connectivity [7-14]. However, the mechanisms by which biological sex influences behavior, and the roles of shared neural circuits in these processes, are largely unknown.

In the nematode *C. elegans*, hermaphrodites (somatically female animals that transiently generate self-sperm) are self-fertile for much of their reproductive lifespan. Males, however, reproduce only by fertilizing hermaphrodites, making the ability to locate mates crucial to their fitness. Because nematodes rely heavily on chemosensation to navigate their environment, pheromones are likely to be especially important to this process. Accordingly, multiple studies have described the ability of hermaphrodite-secreted compounds to attract males [15-19]. The most well-characterized of these are members of the ascaroside family[20], several of which elicit concentration-dependent, sexually dimorphic behavioral responses alone and in combination [17, 18, 21, 22]. In particular, specific concentrations of ascr#3 (also known as C9 and asc-ΔC9) trigger attractive responses in males and, depending on neuromodulatory state, aversion in hermaphrodites [17, 21, 23, 24].

The neural and genetic underpinnings of sex differences in ascaroside-elicited behavior involve multiple mechanisms and are incompletely understood. One mechanism is anatomical: the ascr#3- and ascr#8-sensitive CEM “cephalic companion” sensory neurons are present only in males [25, 26]. Ablation of these four cells reduces ascr#3 attraction, but a substantial sex difference in behavior remains in their absence[17]. Ascaroside detection also involves shared chemosensory circuitry. In both sexes, the ASK amphid neurons detect ascr#3 and promote attraction [17, 23], though again, ASK-ablated males retain some ascr#3 attraction [17, 21]. The influence of CEM and ASK is counteracted by the ADL amphid neurons, which also detect ascr#3 but provoke an aversive response[21]. ADL responds to ascr#3 stimulation in both sexes, but the magnitude of this response is greater in hermaphrodites than males[21]. Despite these findings, the mechanisms by which shared pheromone detection circuits are functionally tuned by genetic sex remain unknown, as does the role of such tuning in generating sexually dimorphic behavior.

Ultimately, all sex differences in *C. elegans* are controlled by chromosomal sex. Through a well-characterized genetic hierarchy, the X-to-autosome ratio determines somatic sex by regulating the autosomal transcription factor TRA-1A[27]. This factor acts as a master regulator: high TRA-1A activity specifies hermaphrodite development and function, while low or absent TRA-1A promotes male development and function. Downstream of TRA-1A, multiple targets control specific sexually differentiated features. Interestingly, several of these targets are members of the DMRT *(doublesex* and *mab-3-related* transcript) family, having conserved roles in the otherwise rapidly evolving mechanisms of animal sex determination [28, 29]. In *C. elegans*, members of this family regulate neurogenesis [8, 30, 31], cell fate specification [32, 33], and circuit connectivity [14, 34]. However, while the physiology of shared circuitry is also modulated by sex in *C. elegans* [16, 21, 35-38], whether DMRT genes contribute to this process is unknown.

Because TRA-1A acts cell-autonomously, cell-type-specific manipulation of its activity can generate sexually mosaic animals, allowing sexually dimorphic phenotypes to be mapped to specific anatomical foci[39]. Here, we use this as an entry point to probe the underpinnings of sexual dimorphism in ascaroside responses, identifying an economical mechanism that couples chromosomal sex to adaptive variation in behavior by tuning the properties of a single pair of shared chemosensory neurons.

## RESULTS

### An ascaroside mixture elicits male-specific, CEM-independent attraction

To measure behavioral responses to ascarosides, we used an ascaroside mixture (ascr#2:ascr#3:ascr#8, 1 μM:1 μM:10 μM) shown previously to potently attract males [17, 18]. We spotted 1 μL drops of this or control solution onto opposing quadrants of a thin bacterial lawn, and ten young adult animals were placed in the center (Fig. 1a). As expected, males preferentially occupied the ascaroside-containing quadrants over the subsequent 90 minutes, while hermaphrodites were modestly repelled from these regions (Fig. 1b, S1a). Solitary males behaved similarly to those tested in groups of ten (Fig. S1b), and we found no evidence that these compounds promoted male aggregation (Fig. S1c). Thus, this assay measures attraction to (or retention by) ascarosides, rather than secondary influences of these compounds on interactions between males.

**Figure 1.**
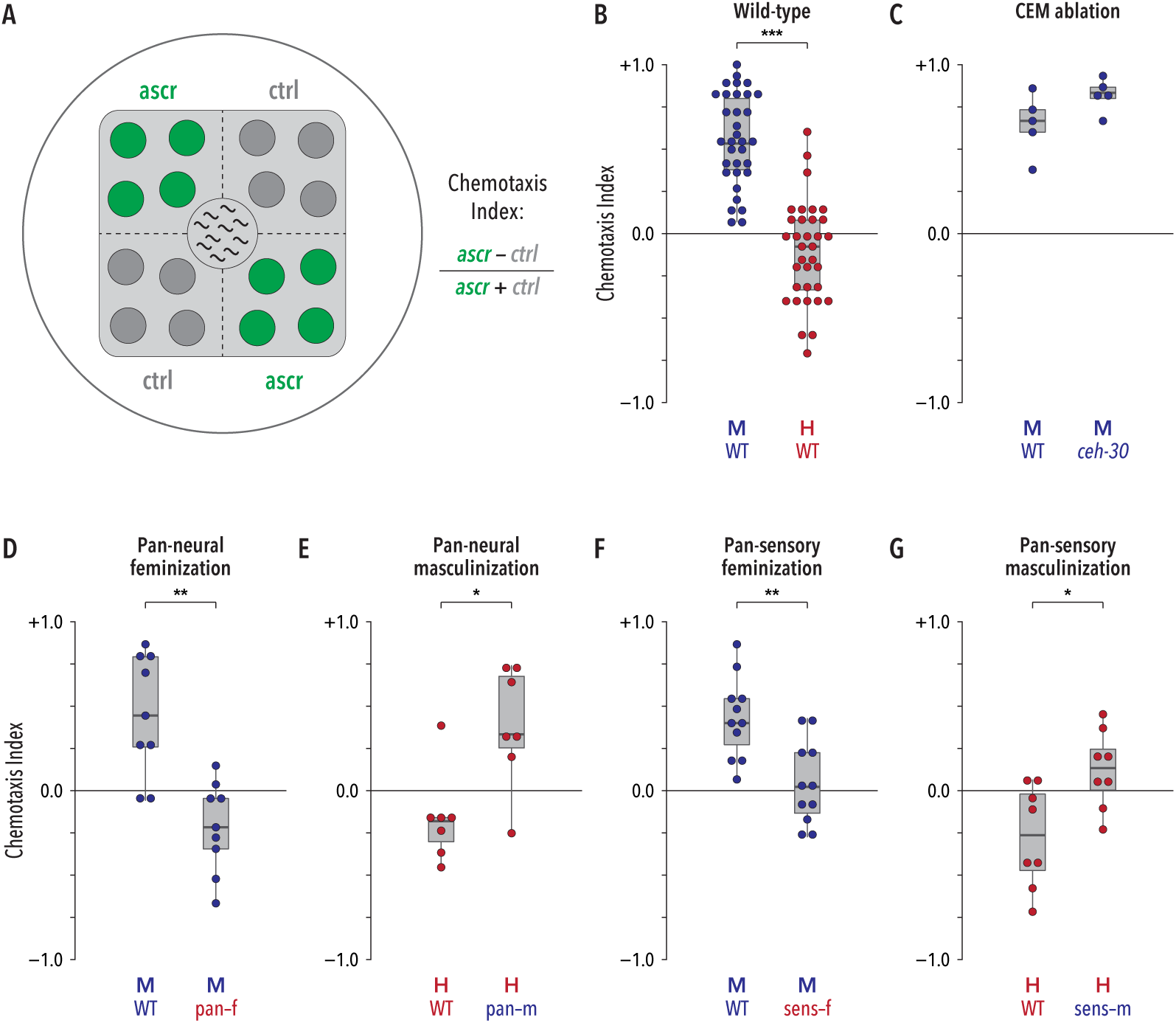
Sexual modulation of shared circuits generates sexually dimorphic behavioral responses to ascarosides. (A) Ascaroside response was measured using a modified quadrant assay in which 1-μL drops of ascaroside or control solutions (green and dark gray, respectively) were spotted on opposite quadrants of a standard culture plate with a square lawn of bacterial food (light gray). Ten animals were placed at the center of the plate and their positions were scored at regular intervals to derive a Chemotaxis Index (see Methods for details). (B) Wild-type animals showed a marked sex difference in behavioral responses to the ascr#2/#3/#8 mixture. (C) Loss of CEM neurons (*ceh-30* mutants) did not detectably influence male behavior. (D, E) Pan-neural feminization (“pan-f”) eliminated ascaroside attraction in males, while pan-neural masculinization (“pan-m”) generated marked attraction in hermaphrodites. (F, G) Feminization of all ciliated sensory neurons (“sens-f”) eliminated ascaroside attraction in males. Masculinization of ciliated sensory neurons (“sens-m”) generated weak attraction in hermaphrodites.

To evaluate the role of the male-specific CEM sensory neurons, we used *ceh-30(lf)* mutants, in which these neurons fail to develop [40, 41]. Unexpectedly, this revealed no detectable impairment in ascaroside attraction (Fig. 1c), suggesting that the roles of different pheromone-detecting neurons depend on behavioral context[17]. As the CEMs are the only sex-specific sensory neurons in the *C. elegans* head, this result also suggests that behavior in this assay might depend on sex-specific modulation of shared circuits.

### Sexual modulation of shared sensory neurons regulates pheromone attraction

To assess the potential for such modulation, we manipulated the sex-determination pathway in post-mitotic neurons. In males, pan-neuronal expression of the *tra-1* activator *tra-2(ic)*[42] feminizes the state of the nervous system but leaves male-specific neurons, as well as the rest of the body, intact[36]. Males carrying this pan-neural feminizing transgene exhibited no pheromone attraction; rather, they showed weak repulsion reminiscent of control hermaphrodites (Fig. 1d). The reciprocal manipulation, genetic masculinization of the hermaphrodite nervous system, can be achieved by pan-neuronal expression of the *tra-1* inhibitor*fem-3* [16, 35]. This too led to a marked change in behavior: masculinized hermaphrodites were attracted to ascarosides, behaving similarly to control males (Fig. 1e). Thus, sexual modulation of shared neurons has a central role in generating sex-typical ascaroside attraction.

We hypothesized that the sex difference in pheromone-evoked behavior might rely on the modulation of sensory neurons. Using a subtype-specific promoter to sex-reverse all ciliated sensory neurons[43], we obtained results qualitatively similar to those above: pheromone attraction was eliminated in males and elicited in hermaphrodites (Fig. 1f, g). The magnitude of these effects appeared smaller than those seen in pan-neural sex-reversed strains; this could reflect differences in transgene expression or might indicate an influence of other sex-specific features. Regardless, these findings indicate a key role for sensory modulation in generating sexually dimorphic pheromone attraction.

### The male state of ADF promotes ascaroside attraction

We next targeted specific sensory neuron classes for sex-reversal. Despite clear roles for the ASK and ADL neurons in behavioral responses to ascr#3[17, 21, 23], specific feminization of these neurons had only modest effects on ascaroside attraction in the quadrant assay. ASK feminization caused a statistically significant but small reduction in attraction, while ADL feminization resulted in a non-significant trend of similar magnitude (Fig. 2a). These results do not rule out important roles for these neurons; rather, they suggest that any such roles are largely independent of genetic sex (see Discussion). We also feminized ASI and ASJ, as these neurons have known or suggested roles in pheromone detection in other contexts [23, 44]. However, neither manipulation had a detectable effect on ascaroside attraction (Figs. 2b, c).

**Figure 2.**
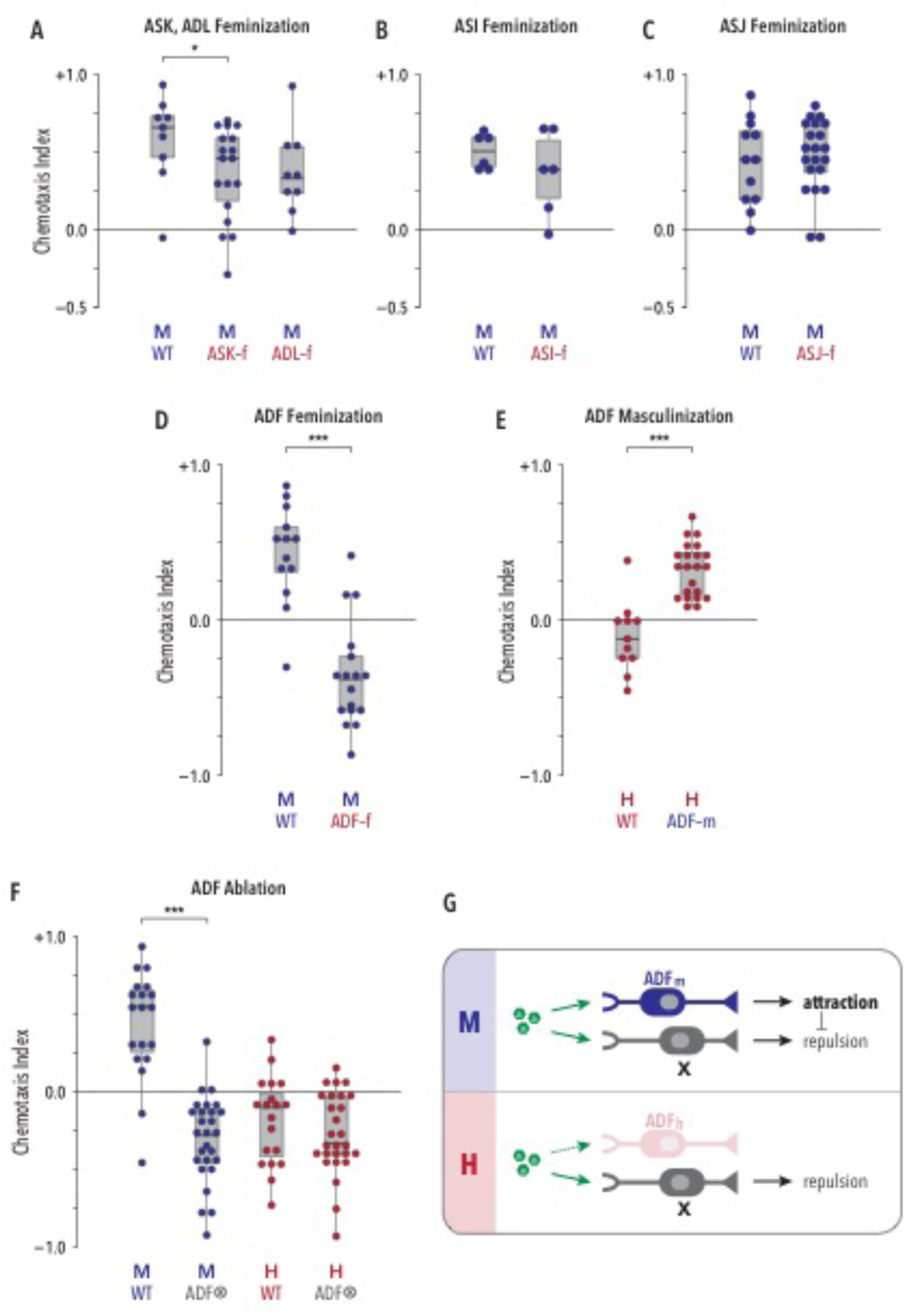
Sex-specific characteristics of the ADF sensory neuron pair determine the valence of the behavioral response to ascaroside pheromones. (A) Genetic feminization of the ASK neurons (“ASK-f”) or the ADL neurons (“ADL-f”) modestly reduced ascaroside attraction in males; this effect was statistically significant only for ASK-f. (B, C) Feminization of the ASI neurons (“ASI-f”) or the ASJ neurons (”ASJ-f”) had no detectable effect on behavior. (D, E) ADF-specific feminization (“ADF-f”) was sufficient to generate pheromone repulsion in males, while ADF masculinization elicited ascaroside attraction in hermaphrodites. (F) Genetic ablation of ADF (ADF0) eliminated sex differences in pheromone response behavior, revealing repulsion in both sexes. (G) ADF sex determines behavioral valence by promoting pheromone attraction in males. In both sexes, an unknown neuronal activity, neuron X (likely ADL[21]; see Discussion), promotes repulsion from ascarosides (green hexagons). In its male state (ADF_m_; upper panel), ADF’s ascaroside-dependent activity overcomes this repulsion to generate attraction. In its hermaphrodite state (ADF_f_; lower panel), ADF is insensitive to ascarosides or is not coupled to attraction circuitry.

In contrast, feminization of the ADF chemosensory neurons led to a pronounced change in behavior: ADF-feminized males, like wild-type hermaphrodites, were repelled from the ascaroside-containing quadrants (Fig. 2d). Conversely, masculinization of ADF in otherwise wild-type hermaphrodites was sufficient to generate marked pheromone attraction (Fig. 2e). Furthermore, the attraction seen in pan-neural masculinized hermaphrodites was suppressed by specifically blocking masculinization in ADF (Fig. S2). These results indicate that male-typical characteristics of ADF are necessary for pheromone attraction and, moreover, that ADF’s sexual state specifies the valence of the behavioral response elicited by pheromones. The ability of ADF masculinization to enable pheromone attraction in hermaphrodites strongly suggests that this behavior requires no sex differences postsynaptic to ADF; that is, ADF signaling likely engages shared navigational circuits to generate sex-specific behavior.

These observations are consistent with two alternative mechanisms. In the first, the male state of ADF would enable pheromone attraction, counteracting a non-sex-specific repulsive influence of other sensory neuron(s). In the second, the hermaphrodite state of ADF would actively promote repulsion, allowing a non-sex-specific attractive drive to dominate only in males. To distinguish between these, we used a split-caspase strategy to ablate ADF [45, 46]. This eliminated attraction in males, revealing an underlying repulsive drive. In contrast, it had minimal effects in hermaphrodites (Fig. 2f). Thus, the male ADF promotes ascaroside attraction, and a non-sex-specific influence promotes repulsion (Fig. 2g). Notably, ADF ablation eliminated the sex difference in behavior (Fig. 2f), demonstrating that its sexual state instructively determines behavioral valence.

### ADF’s genetic sex determines its ability to detect ascarosides

To understand how sexual state influences ADF’s response to the ascaroside mixture, we measured GCaMP3 fluorescence in worms exposed to pheromone stimuli in a microfluidic arena [47, 48]. Stimulation of males with the ascr#2/#3/#8 mixture elicited robust activation of ADF (Fig. 3a), indicating that ADF might directly detect ascarosides. In contrast, ADF appeared completely unresponsive to ascr#2/#3/#8 stimulation in hermaphrodites (Fig. 3b), demonstrating that genetic sex modulates ADF’s sensitivity to ascaroside stimulation. Consistent with this, ADF-specific feminization significantly reduced its responsiveness to ascaroside stimulation: 10 of the 14 ADF-feminized males tested exhibited weak or undetectable calcium responses, while only four of the 22 control males tested failed to respond (Fig. 3a). Furthermore, we observed strong ascaroside-induced calcium transients in most of the ADF-masculinized hermaphrodites tested (Fig. 3b), indicating that “maleness” is sufficient to render this neuron ascaroside-responsive in hermaphrodites.

**Figure 3.**
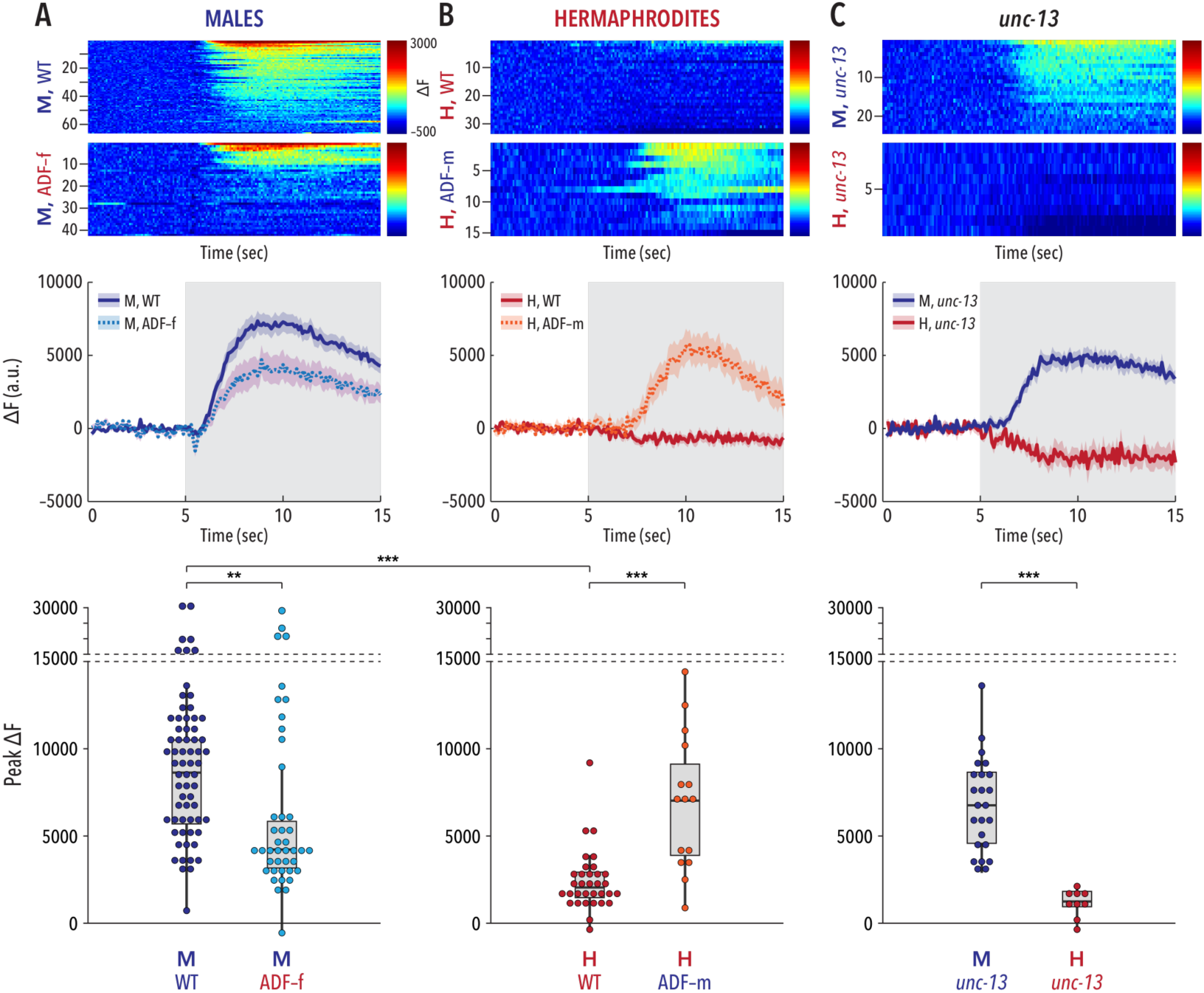
The sexual state of ADF determines its ability to respond to ascaroside stimulation. (A, B, C) *Top panels* show pseudocolored graphs of calcium recordings of individual neuronal responses to stimulation with an ascr#2/#3/#8 mixture. Each animal underwent three rounds of stimulation; each row depicts a single response, sorted by peak activity. The pseudocolor scale represents GCaMP3 ΔF (blue lowest, red highest). *Center panels* show average ΔF intensities ± SEM for the recordings depicted in the upper panels. Gray shading indicates the period of pulsed ascaroside stimulation. *Bottom panels* show peak ΔF values for each recording; dots represent individual responses to each pulse. (A) Wild-type males showed robust calcium transients in ADF upon ascaroside stimulation. The strength of this response was significantly reduced in ADF-feminized males. The modest reduction in mean response likely reflects a bimodal distribution of responses (most clearly visible in the lower panel) arising from variable effects of the ADF feminization transgene. (B) In wild-type hermaphrodites, ADF appeared insensitive to ascaroside stimulation. Masculinization of ADF in an otherwise wild-type hermaphrodite was sufficient to confer ascaroside sensitivity to ADF. (C) *unc-13* mutants maintained a marked sex difference in the response of ADF to ascaroside stimulation. *n* = 9-66 trials (3-22 animals with 3 trials per animal).

To ask whether ADF detects ascarosides directly, we imaged activity in *unc-13* mutants, in which chemical synapses are disabled[49]. These animals exhibited strong, sexually dimorphic responses of ADF to ascaroside stimulation (Fig. 3c). While we cannot rule out the possibility that ascaroside signals are transmitted from other neurons to ADF through gap junctions, electrical connectivity to ADF is sparse and features no obvious male-specific synapses[50] (wormwiring.org). Therefore, our results indicate that genetic sex modulates the chemosensory tuning of ADF, allowing it to detect ascarosides only in males.

We also recorded ADF’s response to individual ascarosides in males. Stimulation with ascr#2 or ascr#8 alone generated only weak responses, but stimulation with ascr#3 alone elicited calcium transients comparable to those seen with the ascr#2/#3/#8 mixture (Fig. S3). Consistent with this, male behavior in quadrant assays using ascr#3 alone was similar to that seen with the ascr#2/#3/#8 mixture (Fig. 4a). Thus, modulation of ADF’s ability to detect ascr#3 likely accounts for sex differences in response to the ascr#2/#3/#8 mixture.

**Figure 4.**
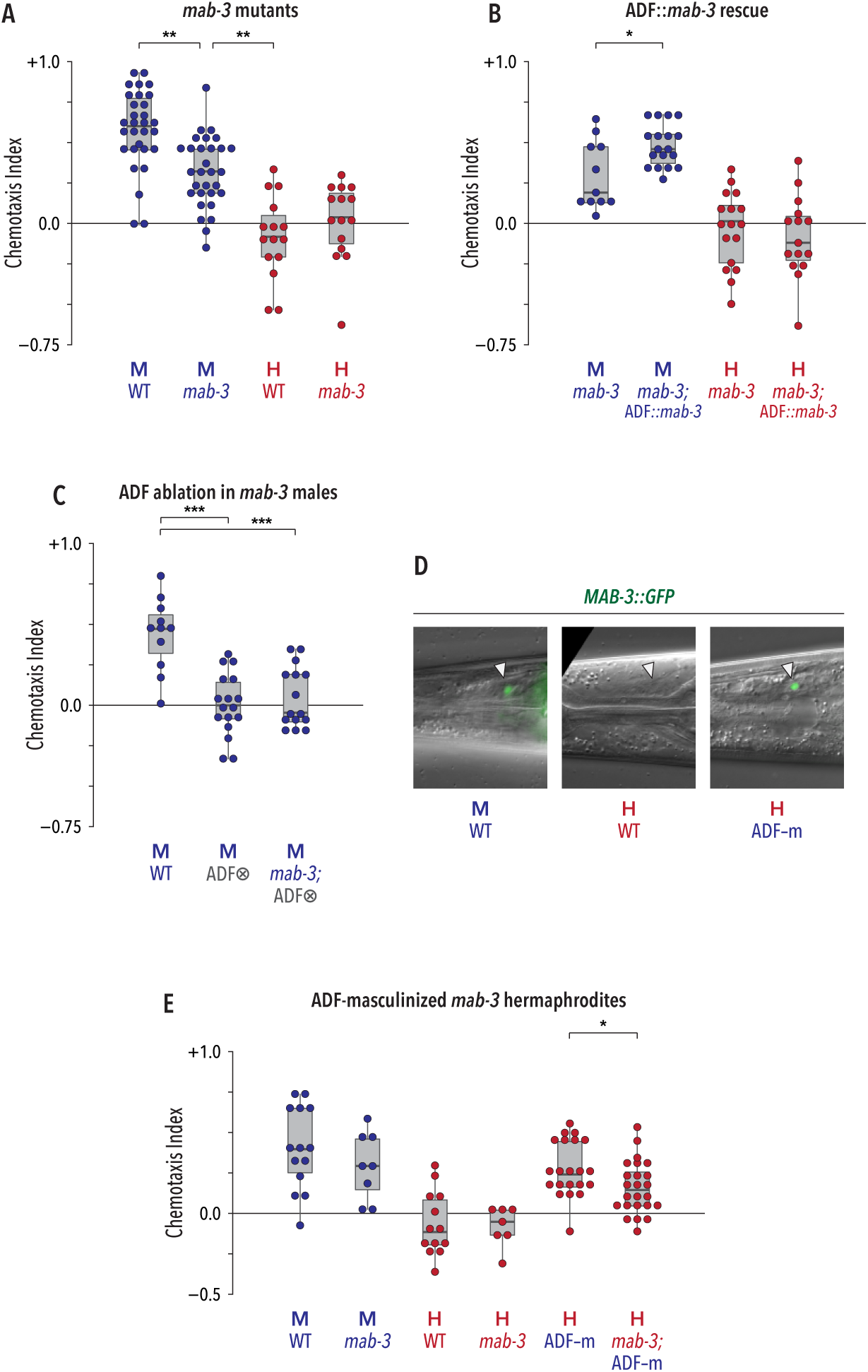
The *doublesex* ortholog *mab-3* acts cell-autonomously and downstream of TRA-1A to modulate ADF function. (A) *mab-3* mutant males had a reduced preference for ascr#3 compared to WT males. Loss of *mab-3* had no detectable effect on hermaphrodite behavior. (B) ADF-specific expression of *mab-3(+)* was sufficient to rescue the behavioral defect of *mab-3* males, but had no noticeable effect on hermaphrodites. (C) Loss of *mab-3* did not further disrupt ascr#3 response when ADF is ablated (ADF⊗). (D) *MAB-3::GFP* was expressed in ADF (white triangles) in WT adult males (*left*) but not in WT adult hermaphrodites (*center*). Genetic feminization of ADF was sufficient to activate ADF in hermaphrodites (*right*). (E) *mab-3* function was required for the full effect of ADF-masculinization on hermaphrodite behavior.

A straightforward explanation for the sexually dimorphic ability of ADF to detect ascr#3 could be the differential expression of an ascaroside chemoreceptor. Of the eight known or putative ascaroside receptors in *C. elegans (srbc-64* and *srbc-66*[51], *srg-36* and *srg-37*[52], *daf-37* and *daf-38*[53], and *srx-43* and *srx-44* [54, 55]), none are known to be expressed in ADF in either sex, making these unlikely candidates. Another putative chemoreceptor, *srd-1*, is expressed male-specifically in the ADF neuron[56], but we observed no pheromone attraction defects in *srd-l(ehl)* null mutants (Fig. S4a). Thus, it remains unclear whether differential receptor expression underlies the sex difference in ADF physiology.

Because ADF is one of the few serotonergic neurons in *C. elegans*, we asked whether serotonin was important for ADF-mediated pheromone attraction. However, males lacking *tph-1* (tryptophan hydroxylase, necessary for serotonin biosynthesis) exhibited wild-type pheromone attraction (Fig. S4b). Thus, ADF likely uses other mechanisms to communicate pheromone detection to downstream circuits.

### The DMRT gene *mab-3* links genetic sex to modulation of ADF function

To understand how ADF’s sexual state modulates its ascaroside sensitivity, we considered known targets of *tra-1*. One of these, *mab-3*, is a founding member of the conserved DMRT or DM-domain family of transcriptional regulators[57]. These genes have roles in sex determination and sexual differentiation in a wide variety of invertebrate and vertebrate species [29, 58]. In *C. elegans, mab-3* implements male-specific features of neurogenesis, intestinal physiology, and tail tip morphology[8, 30, 59]; it also promotes male mate-searching drive[31]. However, a role for *mab-3* in the modulation of neural circuit function has not been noted; indeed, such a role for DMRT genes in any species is not well described.

Interestingly, the only described site of *mab-3* expression among shared neurons is ADF, where its expression is male-specific[31]. We found that *mab-3* mutant males exhibited significantly reduced attraction to ascr#3 and to the ascr#2/#3/#8 mixture, while *mab-3* hermaphrodites appeared wild-type (Fig. 4a, Fig. S4c). Expression of *mab-3(+)* under the control of an ADF-specific promoter rescued the reduced ascaroside attraction of *mab-3* males (Fig. 4b), strongly suggesting that this gene acts cell-autonomously to promote ADF’s male-specific function. Consistent with this, loss of *mab-3* had no further effect on behavior when ADF was genetically ablated in males (Fig. 4c).

To confirm that *mab-3* functions downstream of *tra-1* in ADF, we examined *MAB-3::GFP* expression. As expected, GFP was detectible in ADF in males but not in hermaphrodites; furthermore, masculinization of ADF was sufficient to activate *MAB-3::GFP* in this cell (Fig. 4d). *mab-3* was also partially required for the ascr#3 attraction generated by ADF masculinization in hermaphrodites (Fig. 4e). Notably, however, loss of *mab-3* did not completely eliminate ascaroside attraction in males or in ADF-masculinized hermaphrodites (Figs. 4a, e). In addition, ectopic expression of *mab-3* in the hermaphrodite ADF was insufficient to generate attraction (Fig. 4b). Therefore, *tra-1* likely regulates additional factors that, together with *mab-3*, bring about the fully masculinized state of the male ADF. Nevertheless, these findings establish a clear role for *mab-3* in the modulation of ADF function.

### Sex-specific tuning of ADF adaptively facilitates mate-searching

Our results using synthetic ascarosides suggest that sexual modulation of ADF allows males to use ascarosides as mate-location cues. However, the ascaroside concentrations that males would need to recognize for this purpose [22, 60] are likely to be significantly lower than those used here, where small drops of 1 μM ascr#3 are expected to diffuse to the mid nanomolar range. Furthermore, hermaphrodites secrete complex mixtures of ascarosides (and other compounds) whose composition varies according to multiple factors [61-63]. Indeed, whether *C. elegans* males are capable of locating mates using the ascarosides they produce has, to our knowledge, not been reported.

To explore these issues, we asked whether wild-type males could distinguish between wild-type hermaphrodites and *daf-22* mutants, which lack the ability to synthesize short-chain ascarosides including ascr#2, ascr#3, ascr#8 [17, 18, 64, 65]. In these experiments, we adapted the quadrant assay to use genetically immobilized *(unc-13)* hermaphrodites, rather than synthetic ascarosides, as stimuli (Fig. 5a). We found that wild-type males robustly chose to associate with *unc-13* over *unc-13; daf-22* hermaphrodites (Fig. 5b), clearly demonstrating that males do indeed rely on hermaphrodite ascaroside production to identify potential mates. Wild-type hermaphrodites displayed no such preference, distributing themselves equally among *unc-13* and *unc-13; daf-22* hermaphrodites (Fig. 5b). Furthermore, the ability of males to detect ascarosides in this context depended heavily on the sexual modulation of ADF: feminization of ADF significantly reduced the preference of males for associating with *unc-13* over *unc-13; daf-22* hermaphrodites. Moreover, ADF ablation rendered males barely able to distinguish between the two populations (Fig. 5b). Together, these findings establish a central role for ADF in males’ ability to respond to ascaroside stimuli encountered in an ethologically relevant context.

**Figure 5.**
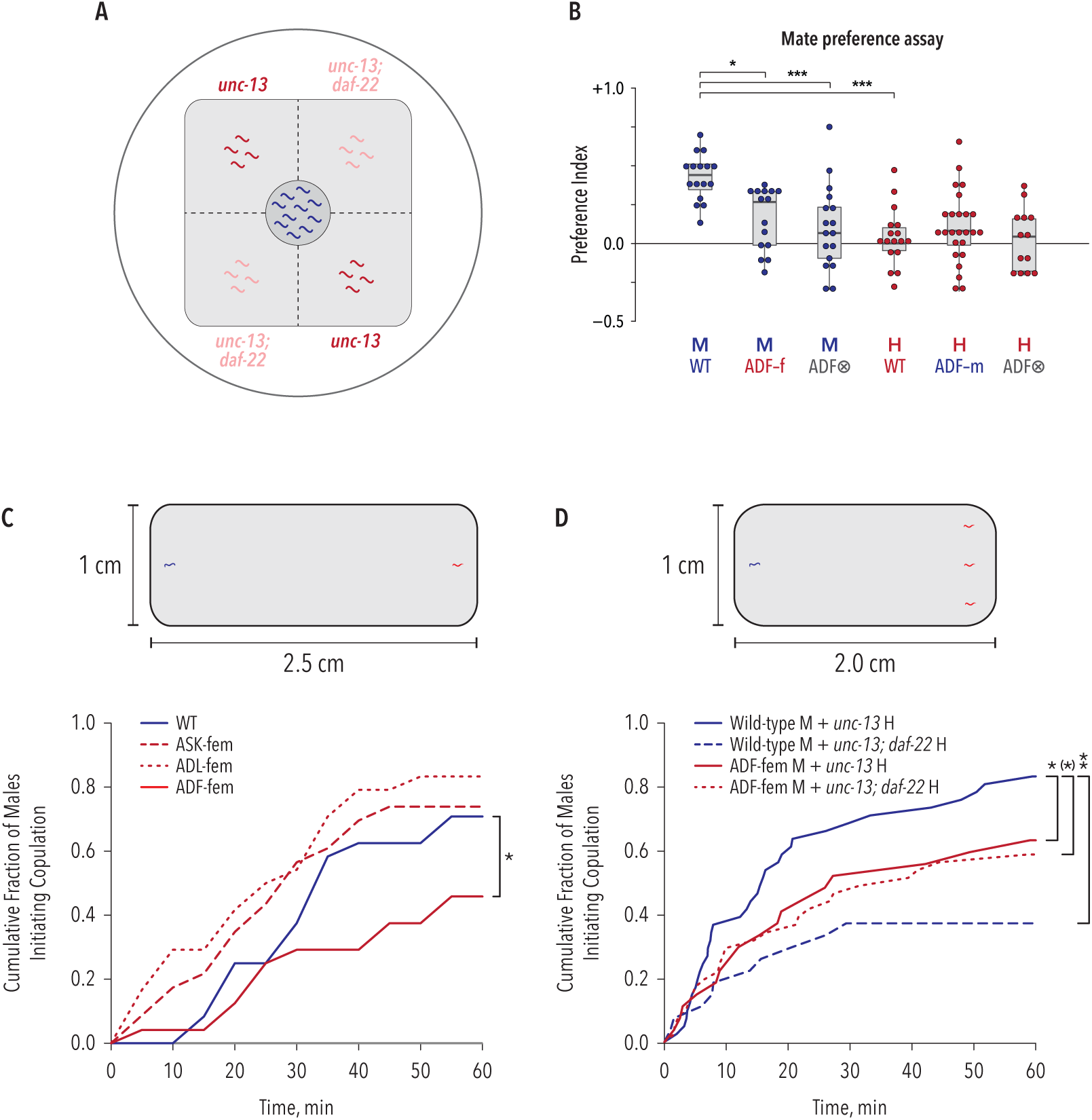
Male-specific tuning of ADF is necessary for males to use ascarosides to locate mates. (A) Modified quadrant assay designed to measure the ability of males to distinguish between ascaroside-producing (*unc-13*) hermaphrodites and ascaroside-lacking (*unc-13; daf-22*) hermaphrodites. (B) Wild-type males exhibited a marked preference for associating with ascaroside-producing over ascaroside-lacking hermaphrodites, while wild-type hermaphrodites showed no bias toward either. ADF feminization (ADF-f) reduced this preference in males, and ADF ablation (ADF⊗) nearly eliminated it. Neither ADF masculinization (ADF-m) nor ADF⊗ had measurable effects in hermaphrodites. (C, D) *Upper*, diagrams of assay conditions. Gray regions represent lawns of *E. coli*. The starting points of males (blue) and hermaphrodites (red) are shown. *Lower*, cumulative rates with which males locate and initiate copulation with hermaphrodites. (C) Feminization of ADF compromised the ability of males to locate and copulate with hermaphrodites, while feminization of ASK or ADL had no detectable effect. (D) Wild-type males took longer to find and copulate with *daf-22* hermaphrodites (dashed blue line) compared to wild-type hermaphrodites (solid blue line). Feminization of ADF (solid red line) impaired males' ability to find mates; it also eliminated the ability of males to use hermaphrodite-produced ascarosides as mate-location cues (compare solid red to dashed red lines).

In hermaphrodites, neither ADF masculinization nor ADF ablation brought about a detectable preference of hermaphrodites for *unc-13* over *unc-13; daf-22* hermaphrodites (Fig. 5b). Thus, “maleness” in ADF alone appears insufficient alter hermaphrodite behavior in this context, suggesting roles for other sensory neurons—either shared or male-specific—in generating male-typical behavior in this setting.

To better understand the effects of hermaphrodite-produced ascarosides on male behavior, we placed single males on a small food lawn 2.5 cm away from a single *unc-13* adult hermaphrodite (Fig. 5c) and monitored their position and behavior at five-minute intervals. Under these conditions, roughly 70% of control males located the hermaphrodite and initiated copulatory behavior within 60 min (Fig. 5c). ADL- and ASK-feminized males behaved comparably to wild-type, but ADF-feminized males were significantly impaired in this task, with fewer than 50% initiating copulation within the same time frame (Fig. 5c). This is not likely to be secondary to other causes, as we found no difference in exploratory activity between control and ADF-feminized males (Fig. S5a) and ADF-feminized males exhibited no significant reduction in their propensity to initiate mating once hermaphrodite contact occurred (Fig. S5b). These results suggest that the male state of ADF allows males to use hermaphrodite-derived ascarosides as navigational cues.

We next asked whether ascaroside production was necessary for the ability of ADF to promote mate-finding. In these experiments, we placed single males 3 cm from a group of three hermaphrodites (Fig. 5d) and scored male behavior using recorded video. We found that males took significantly longer to initiate copulation with *unc-13; daf-22* hermaphrodites compared to control *unc-13* hermaphrodites (Fig. 5d). This is not explained by a defect in copulatory behavior itself: male contact-response efficiency was not compromised by a loss of ascaroside production by its mate (Fig. S5c). The ability of ADF-feminized males to locate mates and initiate copulation, already reduced compared to WT animals, was not further degraded by pairing them with *daf-22* mutant hermaphrodites (Fig. 5d). Thus, ADF facilitates the ability of males to use hermaphrodite-derived ascarosides as navigational cues. Unexpectedly, WT males appeared to locate *daf-22* hermaphrodites even less efficiently than did ADF-feminized males, though this difference was only marginally significant (*p* = 0.093) (Fig. 5d). If biologically meaningful, this result might suggest that ADF feminization triggers the development of compensatory, ascaroside-independent sensory mechanisms.

## DISCUSSION

Pheromone attraction provides an ideal entry point for understanding the genetic, developmental, and physiological mechanisms that bring about naturally occurring, adaptive behavioral variation[66]. Here, we find that ADF, a pair of head chemosensory neurons present in both sexes, is a central focus of the neuromodulatory activities that generate sex differences in *C. elegans* pheromone attraction. ADF detects the ascaroside ascr#3 only in males, a result of cell-autonomous regulation by terminal sexual regulator *tra-1* and its direct target *mab-3*, a conserved DMRT gene [29, 58]. This chemosensory switch determines the nature of an animal’s behavioral response to ascarosides and allows males to use the trace amounts of ascarosides produced by individual hermaphrodites as navigational cues that facilitate mate-searching. Together, this reveals a mechanism by which genetic sex acts through a conserved transcriptional regulator to focus male sensory function on a stimulus with high salience for male reproductive fitness (Fig. 6).

**Figure 6.**
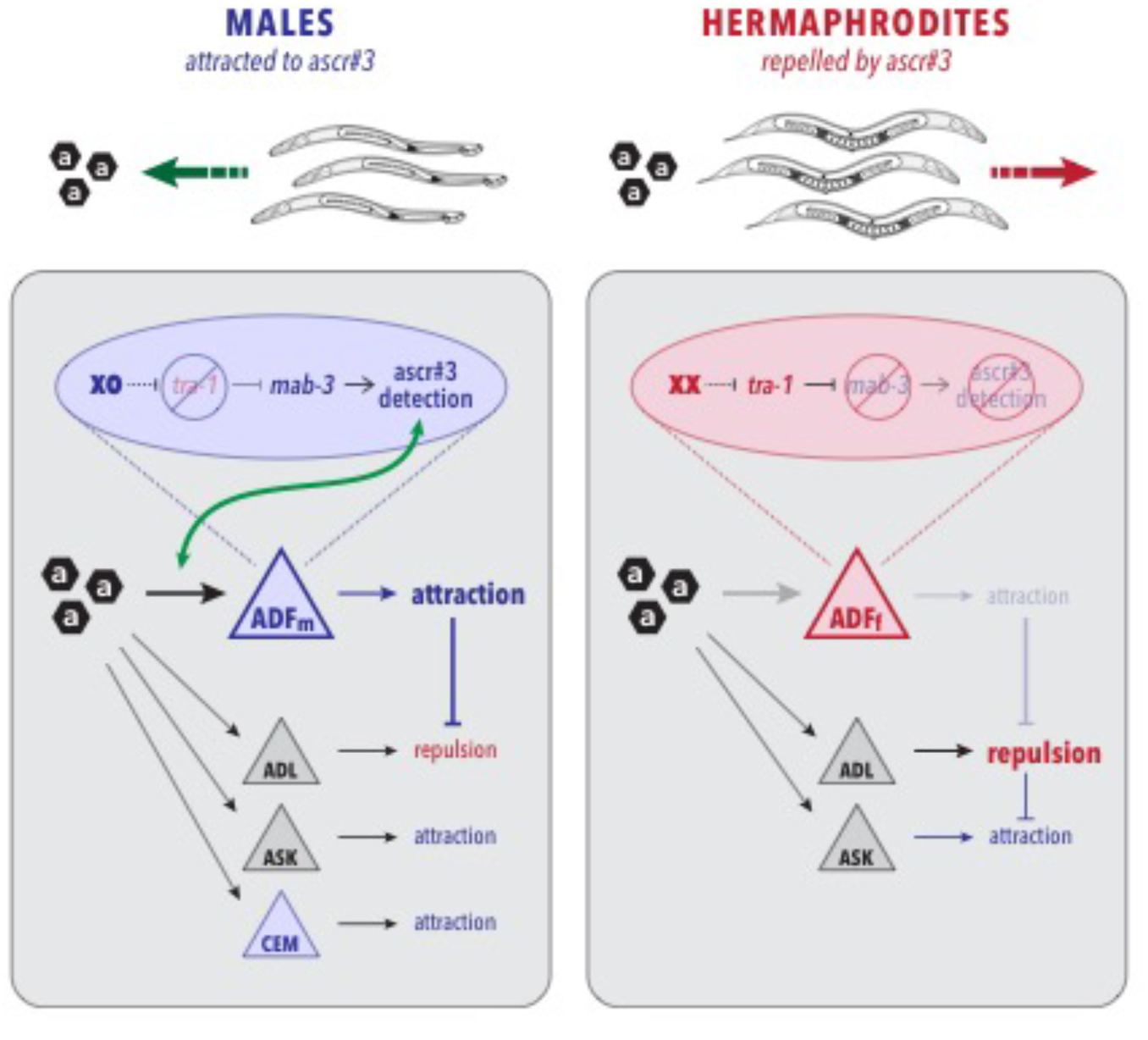
Cell-autonomous modulation of ADF sensory function by genetic sex is sufficient to determine behavioral valence. Black hexagons depict ascarosides; triangles are sensory neurons. Blue triangles indicate the male state of the shared neuron ADF (ADF_m_) and the male-specific CEM neurons. The red triangle indicates the hermaphrodite state of ADF (ADF_h_). Gray triangles depict shared neurons in which genetic sex does not appear to have a prominent impact on function. The blue and red ovals show the genetic mechanism that links chromosomal sex to ascaroside detection by ADF in its male and hermaphrodite states, respectively. In males, genetic sex acts through *mab-3* to confer the ability to detect ascarosides onto ADF, which in turn engages shared circuitry to promote attraction. This drive overcomes a non-sex-specific repulsion from ascr#3, likely generated by ADL. The ASK and CEM neurons also promote ascaroside attraction, though their roles in the assay used here appears to be secondary. In hermaphrodites, *tra-1* represses *mab-3*, preventing ADF from detecting ascaroside stimuli. This likely allows ADL-dependent repulsion to predominate over any attractive drive that might be provided by ASK.

While early experiments suggested that ADF may have a role in the detection of dauer pheromone[44], a role for the ADF neurons in sex pheromone attraction has not been previously described. In hermaphrodites, ADF is best known for its roles in detection of food-related compounds, release of serotonin, inhibition of entry into the dauer stage, and modulation of food-dependent behaviors [44, 67-70]. Our results indicate that, in males, ADF also detects ascr#3 (and perhaps other ascarosides), and that the transmission of this signal is independent of serotonin release. An alternative model, that ADF simply provides a food signal permissive for pheromone attraction, seems unlikely, as such a signal would be expected to be serotonin-dependent. Furthermore, that male-specific properties of ADF are important for pheromone attraction is difficult to reconcile with such a model, as there are no prominent sex differences in ADF’s synaptic output (wormwiring.org). Furthermore, ADF responds to ascaroside stimulation with a calcium transient even when chemical synaptic transmission is disrupted, strongly suggesting that ADF itself is a detector of the pheromone signal. We cannot rule out the possibility that the male ADF receives ascaroside signals indirectly via gap junctions; however, it is again difficult to see how such a mechanism could work, as connectomics studies have revealed no male-specific electrical synapses involving ADF (wormwiring.org).

Three additional *C. elegans* sensory neuron types—CEM, ASK, and ADL—also have important roles in ascaroside-elicited behaviors[17, 21, 23] (Fig. 6). The CEMs, the only male-specific sensory neurons in the worm’s head[26], respond to ascaroside stimulation with a complex pattern of activity that is thought to facilitate the detection of changes in ascaroside concentration[25]. In a pheromone-attraction assay distinct from the one used here, CEM ablation reduces, but does not eliminate, attraction[17]. Why our assay reveals no effect of CEM loss is unclear, but likely reflects dependencies on the concentration and diffusion of the stimulus as well as the specific behavioral responses being measured. The ASK neurons respond to ascaroside stimulation in both sexes[23], explaining why a prominent role for sexual modulation of ASK was not revealed by our studies. (We did observe a small but statistically significant decrease in the ascaroside attraction of ASK-feminized males, suggesting that this neuron could indeed have sex-specific properties; under different conditions, these might have an even more pronounced role in male behavior.) The ADF neurons, unlike the CEMs and ASK, promote aversion to high concentrations of ascr#3 in hermaphrodites. ADL detects ascarosides in both sexes, but its response is blunted in males[21]. This makes ADL a strong candidate for mediating the underlying repulsive behavior revealed by feminizing or ablating of ADF in males, indicated by neuron “X” in Fig. 2g.

Our findings indicate that the DMRT transcription factor *mab-3* couples chromosomal sex to the modulation of ADF function. In *C. elegans*, DMRT genes have been implicated in male-specific morphogenesis, neurogenesis, and circuit connectivity [8, 14, 30-32, 34, 59]. In Drosophila, the DMRT gene *doublesex* has important functions in neurogenesis and cell death, but circuit connectivity and behavior are largely controlled by the insect-specific transcription factor *fruitless*[71]. In neither of these systems, however, have DMRT genes been found to directly modulate the functional characteristics of sexually isomorphic circuits. DMRT genes also act in many other organisms to control sex determination and sexual differentiation [29, 58], but little is known of their potential functions in the nervous system. Consistent with the male-specific expression of *mab-3* in ADF[31], we found that this gene acts cell-autonomously to promote ascaroside attraction, and that *mab-3* is required for the full effects of ADF masculinization on hermaphrodite behavior. Thus, *mab-3* couples several distinct aspects of male development and physiology downstream of the sex determination hierarchy. Moreover, these findings show that DMRT genes can modulate the physiological properties of shared neurons to influence circuit function and behavior, further solidifying their central role in sculpting sex-specific features of the invertebrate nervous system. Because *mab-3* expression in hermaphrodites appears insufficient to generate pheromone attraction, and because *mab-3* loss in males only partially compromises pheromone attraction, it is likely that other factor(s) act with *mab-3* to fully masculinize ADF. Identifying these factors, as well as their downstream targets, will be an important focus of future work.

Because ADF masculinization is sufficient to sensitize this neuron to ascarosides and to generate an attractive response to them in hermaphrodites, no sex-specific features post-synaptic to ADF seem to be necessary for ascr#3 attraction. An appealing idea is that male-specific ascaroside detection results from the regulation of ascaroside chemoreceptor(s) by genetic sex. However, since none of the known ascaroside receptors appear to be involved ADF-mediated attraction, testing this model will require further studies. Regardless, our results reinforce the idea that sex differences in behavior can emerge not only via sex-specific circuits but also through the differential activation of programs common to both sexes [72-74]. Here, we favor the idea that ascaroside detection by the male ADF differentially engages shared navigational circuits. Indeed, ADF activation is associated with locomotor slowing in hermaphrodites [68, 70] (though ADF-derived serotonin is also implicated in promoting roaming[75]), and exposure to ascr#3-containing plates increases male reversal rate[17]. Therefore, sex-specific detection of ascarosides by ADF might trigger a dwelling or local-search like state that enables males to more thoroughly investigate their immediate surroundings.

In insects, classic and recent studies have found that sex differences in behavioral responses to sex pheromones emerge through multiple mechanisms. Significant attention has been paid to the role of the sex-specific processing of chemosensory information, largely as a result of sexually dimorphic circuits[9, 12, 76]. Sex differences in the number and connectivity of sensory structures also have important roles [77, 78]. Interestingly, however, like in *C. elegans*, insect chemosensory structures can also be sex-specifically tuned[79]. Thus, simply blinding one sex to a stimulus may be an efficient and conserved mechanism for generating sex-specific attention.

Sex-specific tuning of sensory function is emerging as a general mechanism for generating sex differences in *C. elegans* behavioral prioritization and decision making. The sexually modulated function of ADF has clear parallels in another chemosensory neuron, AWA, also a target of cell-autonomous regulation by genetic sex[37]. In AWA, regulated expression of the chemoreceptor ODR-10 mediates sex differences in the attraction to the food-related cue diacetyl and to bacterial food itself, allowing well-fed adult males to prioritize mate-searching over feeding[37]. Sex differences in sensory function are also implicated in male responses to unknown, non-ascaroside components of hermaphrodite-conditioned media[16]. The shared chemosensory neuron ASJ is also modulated by sex, regulating its production of DAF-7/TGFß and influencing male mate-searching behavior, though whether this reflects a sex difference in ASJ sensory function *per se* is unclear[38]. The degree to which the function of the other 56 shared, ciliated sensory neurons in *C. elegans* is also modulated by sex is unknown, but it could be much deeper than has been appreciated. Notably, many other aspects of *C. elegans* internal state regulate chemosensory neuron function [24, 37, 80-90], suggesting that such plasticity provides a specific, flexible, and economical means for the compact *C. elegans* nervous system to implement state-dependent behaviors. Regulated chemosensation is also emerging as an important mechanism in many other systems, including insects, rodents, and humans [84, 88, 91-96]. Thus, sex differences in chemosensory tuning can be considered instances of more general mechanisms that generate flexibility in sensory circuit function.

A hallmark of nervous systems is their ability to detect and respond to stimuli in a manner commensurate with internal and external conditions. Neuromodulation is central to this state-dependence, generating plasticity in behavior by reconfiguring circuit dynamics [97, 98]. With regard to some aspects of internal state—e.g., hunger, sleep, and aggression [99-101]—significant progress has been made in the role of neuromodulation. Though biological sex is also deeply conserved dimension of internal state, how it influences circuit function is poorly understood, and whether it is useful to consider sex as a modulatory influence is unclear [1, 2, 102, 103]. However, such a view could provide a useful framework for understanding the sex-biased susceptibility seen in many neurological and psychiatric disorders[104]. Continued studies of the mechanisms by which biological sex regulates neuronal function, particularly in the context of sex-shared circuits, are likely to shed light on these issues.

## MATERIALS AND METHODS

### Nematode strains

Nematode culture was carried out as described[105]. Strains used in this study are listed in Table S1. For simplicity, the reference strain *him-5(e1490)* V is referred to here as “wild-type”. Unless otherwise noted, all strains used contained this mutation, which increases the frequency of males in hermaphrodite self-progeny.

### Molecular Biology

Sex-reversal transgenes were generated using the Multisite Gateway Cloning System (Invitrogen). Neuron-specific promoters (amplified with primers shown in Table S2) were used to drive expression of *fem-3(+)* or *tra-2(IC)* to masculinize or feminize, respectively. Transgenic animals were generated using microinjection. Injection mixes contained 100 ng/μl of the co-injection marker *Pelt-2::gfp* and 50 ng/μl of the sex-reversal (or *mab-3* rescue) construct.

### Ascarosides

Synthetic ascarosides[64] were reconstituted in 2mL EtOH and stored at -20°C. Stock ascarosides were diluted to working concentrations with Milli-Q water, aliquoted into single-use tubes, and stored at -80°C.

### Behavior

#### Quadrant Assay

The day before the assay, L4 animals were picked to sex-segregated plates (15 animals per plate) and assay plates were prepared. A custom-made stamp was used to mark quadrant boundaries on the bottom of unseeded 6-cm NGM plates. The quadrant area comprised a 3-cm square divided into four zones with a 0.5-cm radius circle at the center. After spreading 50 μl of *E. coli* OP50 culture within the 3 cm square, plates were incubated at 20°C. The day of the assay, eight 1-μL drops of ascaroside solution (either 1 μM ascr#2, 1 μM ascr#3, 10 μM ascr#8 together or 1 μM ascr#3 alone, as indicated) or control (equivalent amounts of ethanol diluted in water) were placed onto the surface of opposing pairs of quadrants (Fig. 1a). At time t=0, ten worms of the appropriate sex and genotype were placed in the center circle. At t=30 min, t=60 min, and t=90 min, the number of worms in each pair of quadrants was recorded. Chemotaxis indices for the three time points [calculated as CI = (# in ascaroside quadrants - # in control)/(# in ascaroside quadrants + # in control)] were averaged together to generate a final CI value for each assay. Averaging these three points provides a more reliable measure of the ascaroside-response behavior of the population; there was no evidence of adaptation to the stimulus during this time (Fig. S1a). Statistical significance was assessed using the Mann-Whitney-Wilcoxon test or the Kruskal-Wallis test with Bonferroni correction and Dunn’s posthoc test. *p* values are indicated as follows: * *p* < 0.05, ** *p* < 0.01, *** *p* < 0.001.

#### Quadrant Assay with Immobilized Hermaphrodites

These assays were carried out using the same overall format as the *Quadrant Assay* described above. Instead of ascaroside and control solutions, genetically paralyzed *unc-13* or *unc-13; daf-22* hermaphrodites were placed in opposing quadrants (four worms per quadrant). After 30 min, ten worms of the appropriate sex and genotype were placed in the center circle (*t* = 0). Plates on which hermaphrodites moved beyond the boundaries of their quadrant (an uncommon occurance) were censored.

#### Mate Localization Assay (Manual)

The day before the assay, L4 males were picked to individual plates and L4 *unc-13(e51)* hermaphrodites were picked to a single plate. Assay plates were prepared by spreading 10 μl of *E. coli* 0P50 into a 1 cm x 2.5 cm lawn in the center of a 6-cm NGM plate and were incubated at 20°C overnight. Two hours before the start of the assay, one *unc-13* hermaphrodite was placed at one end of the bacterial lawn on each plate. The assay began when an adult male was placed at the opposing end of the lawn. All plates were checked at 5-min intervals for 60 min and the time at which the male was first observed mating with the hermaphrodite was recorded. Mating was defined by the male tail contacting the hermaphrodite and scanning for (or already at) the vulva. The fraction of males observed mating by time *t* was plotted. Kaplan-Meier survival curves were generated and compared using the log-rank test.

#### Mate Localization Assay (Video)

These experiments were performed as described in *Mate Localization Assay (Manual)* above, except that the lawn size was reduced to 1 cm x 2 cm to fit smaller 3.5-cm plates, and 3 *unc-13* or *unc-13; daf-22* hermaphrodites were placed at one the end of the lawn. A Basler acA2500-14um camera connected to a Fujifilm HF16Sa-1 lens was mounted above an illumination source, and Pylon Viewer software was used to capture images of the entire lawn at 1 or 2 frames/sec for 60 min. Interactions were defined as any physical contact between the male and a hermaphrodite. Mating initiation was scored as the first sustained contact that resulted in tail scanning behavior. Kaplan-Meier survival curves were generated and compared using the log-rank test with the Holm correction.

#### Male Mating Behavior

Male response behavior was assayed as described[106]. Contact response efficiency was evaluated using the Mate Localization video data. Response efficiency was calculated as the reciprocal of the number of interactions until mating initiation occurred.

#### Exploratory Behavior

Experiments were performed as described in *Mate Localization Assay (Video)* above except that no hermaphrodites were picked to the assay plate. Trajectories of individual males were tracked using a modified version of the Parallel Worm Tracker Matlab script[107]. Trajectories were passed through a grid of 0.9mm^2^ squares and the number of squares covered by each worm trajectory was determined. Statistical analysis was performed using unpaired Student’s t-tests.

### Neuronal calcium imaging

#### Automated imaging and stimulation

Neuronal imaging was performed using a Zeiss AxioObserver.A1 inverted microscope with Zeiss Plan-Apochromat objective lens (10x/0.45 NA) and a Hamamatsu Orca Flash 4.0 sCMOS camera mounted with a 1.0x C-mount adapter. MicroManager software was used to acquire image stacks (10 frames s^-1^ for 30 s per trial) and to control liquid stimulus delivery (5 -15 s) via a microscope controller (Nobska Imaging) that actuated Parker solenoid valves via a Automate ValveLink 8.2 controller. Excitation illumination pulses (10 ms per frame) were delivered from a 50W blue LED (Mightex) or Lumencor SOLA through an EGFP filter set. For testing individual ascarosides in sequence, the inlet tube was transferred manually to each ascaroside tube and allowed to flow for 1 min between stimuli.

#### Microfluidic device designs

Stimulus pulses were delivered using a modified two-arena microfluidic device as previously described[47], allowing simultaneous recording from two populations. One of two stimulus streams was directed into the arenas using a computer-controlled three-way valve that switched the flow position of a "control" fluid stream, while the other stimulus stream bypassed the arena directly to the outflow. This design allowed chemical stimuli to be changed during an experiment while animals were presented a buffer solution and remained naive to the next stimulus.

#### Microfluidic device fabrication

Reusable monolayer microfluidic devices were prepared using soft lithography as previously described[48]. After each use, devices were cleaned in 95% EtOH at least overnight to remove residual PDMS monomers, rinsed in water and baked for at least 30 min at 65 °C to evaporate any absorbed EtOH. Devices were reversibly sealed against a hydrophobic glass slide with a support glass slide containing inlet and outlet holes drilled with a diamond-coated bit above the PDMS device and clamped in a stage adapter.

High-purity Teflon PFA tubing was used to prevent cross-contamination when presenting multiple stimuli within an experiment and for all inlet valves. Waste and animal loading connections were made with microbore tubing containing a metal tube on one end for insertion into the microfuidic device[48]. Buffers and stimuli were delivered from 30 mL syringe reservoirs, or 1.5 mL tubes when switching between stimuli in the same experiment. Fluids were delivered by hydrostatic pressure, with flowrate controlled by the distance (∼70 cm) between inlet reservoirs and the outlet reservoir.

#### Stimulus preparation

Stimulus dilutions were prepared fresh on the day of the experiment from stock solutions of ascr#2 (1.93 mM), ascr#3 (1.41 mM), ascr#8 (0.64 mM) (all in 100% EtOH and stored at -20°C), 1X S basal kept at 20°C, and 1 M (-)-tetramisole (Sigma) in 1X S basal kept at 4°C. For all imaging experiments, dilutions were made at room temperature in 1 mM (-)-tetramisole 1X S basal buffer to paralyze body wall muscles and keep animals stationary. In experiments using ascaroside mixtures, the final stimulus solution for imaging was a 0.1:0.1:1.0 μM mix of ascr#2:ascr#3:ascr#8 with 0.169% EtOH. Buffer and control solutions also contained 0.169% EtOH. In experiments using individual ascarosides, the stimulus solutions contained each single ascaroside at a final concentration identical to that in the ascaroside mixture with all solutions containing a final 0.169% EtOH concentration.

#### Experimental setup

Microfluidic arenas were assembled and degassed in a vacuum desiccator for at least 30 min before loading buffer through the outlet port. Inlet reservoirs were connected to the arena via microbore tubing[48] and the device was flushed with S basal buffer. After flow switching was verified, animals were gently injected into the arena. (In all imaging experiments, animals were tested as young adults when selected as L4s the day prior and males of a similar size.) Buffer flow continuously washed the animals, and animals were paralyzed with 1 mM tetramisole for ∼1 h before recording and stimulating. After each experiment, devices were disassembled and washed to be reused as previously described[48].

#### Data analysis and statistics

ADF neural fluorescence was analyzed from video frames using a modified ImageJ NeuroTracker software suite and custom MATLAB scripts[47]. Background-corrected integrated neural fluorescence traces *F(t)* were subtracted by baseline fluorescence F_0_ (mean for the first 4 s) to obtain the corrected calcium response Δ*F*) for each animal and stimulation trial. The peak change in neural fluorescence was quantified by finding the max Δ*F* between 1 second after the start and 5 seconds before the end of the stimulus pulse. Data were analyzed using a Kruskal-Wallis test with Bonferroni correction and Dunn’s posthoc test. Heat maps were sorted vertically by the peak change in neural fluorescence across all individuals and repeated trials.

Because the ADF::GCaMP transgene *syEx1249* expressed GCaMP3 at very low levels, baseline fluorescence values (F_0_) were close to zero. Data are reported as raw ΔF values rather than normalized ΔF/F_0_ to avoid excess noise and artifact. To confirm that the baseline GCaMP signal did not differ between strains, we used high-power epifluorescence microscopy to quantitate baseline fluorescence in UR1031 (wild-type background) and UR1053 (ADF-feminized) adult males. GCaMP3 fluorescence was imaged using a 40x Plan-NEOFLUAR objective on a Zeiss Axioplan 2 microscope equipped with a Hamamatsu ORCA ER camera. Fluorescence intensity was quantified using ImageJ version 1.51j as described[108]. Briefly, an outline was drawn around ADF and the area and integrated density was measured. Background fluorescence was obtained by outlining a circular area near ADF and measuring the mean fluorescence. Total Corrected Cellular Fluorescence was determined by TCCF = Integrated Density - (Area x Mean Background Fluorescence). Using this method, we obtained TCCF values for UR1031 and UR1053 of 4.54 ± 2.26 (*n*=13) and 4.91 ± 2.95 (*n*=19), respectively. An unpaired Student’s t-test did not indicate that these values differed significantly (*p*=0.693).

---

**Table S1.** Strain list.

| Strain | Genotype |
| --- | --- |
| PT8 | *pkd-2(sy606) IV; him-5(e1490) V* |
| UR226 | *him-5(e1490) V; fsEx160[Posm-5::fem-3(+):SL2::mCherry, Punc-122::GFP]* |
| UR236 | *him-5(e1490) V; fsIs15[Prab-3::fem-3(+):SL2::mCherry; Punc-122::GFP]* |
| UR278 | *mab-3(e2093) II; him-5(e1490) V* |
| UR754 | *him-5(e1490) V; fsEx357[Posm-5::tra-2(ic):SL2::mCherry]* |
| UR866 | *him-5(e1490) V; fsEx317[Prab-3::tra-2(ic):SL2::mCherry; Punc-122::GFP]* |
| UR926 | *him-5(e1490) V* |
| UR940 | *him-5(e1490) V; fsEx449 [Psrh-142::tra-2(ic):SL2::mCherry; Pelt-2::GFP line 1]* |
| UR945 | *him-5(e1490) V; fsEx454 [Pgpa-4::tra-2(ic):SL2::mCherry; Pelt-2::GFP line 1]* |
| UR948 | *him-5(e1490) V; fsEx457 [Psrbc-64::tra-2(ic):SL2::mCherry; Pelt-2::GFP line 1]* |
| UR949 | *him-5(e1490) V; fsEx458 [Psrbc-64::tra-2(ic):SL2::mCherry; Pelt-2::GFP line 2]* |
| UR963 | *him-5(e1490) V; fsEx461 [Psrh-220::tra-2(ic):SL2::mCherry; Pelt-2::GFP line 1]* |
| UR987 | *him-5(e1490) V; fsEx212[Pelt-2::GFP; Psrh-142::GFP; Psrh-142::ced-3(p15); Psrh-142::ced-3(p17) line 2]* |
| UR999 | *him-5(e1490) V; fsIs15[Prab-3::fem-3(+):outtron::mCherry+Punc-122::GFP]; fsEx449[Psrh-142::tra-2(ic)::mcherry unc-54; Pelt-2::gfp line 1]* |
| UR1004 | *him-5(e1490) V; fsEx478[Psrh-142::fem-3(+):SL2::mCherry; Pelt-2::GFP line 1]* |
| UR1026 | *him-5(e1490) V; fsEx482[Pssu-1::tra-2(ic):SL2::mCherry; Pelt-2::GFP line 1]* |
| UR1031 | *pha-1(e2123) III; him-5(e1490) V; syEx1249[Psrh-142::GCaMP3; pha-1(+)]* |
| UR1034 | *pha-1(e2123) III; him-5(e1490) V; fsEx478[Psrh-142::fem-3::mCherry::SL2::unc-54; Pelt-2::gfp]; syEx1249[Psrh-142::GCaMP3+pha-1(+)]* |
| UR1053 | *pha-1(e2123) III; him-5(e1490) V; fsEx449[Psrh-142::tra-2(ic)::mCherry; Pelt-2::gfp]; syEx1249[Psrh-142::GCaMP3; pha-1(+)]* |
| UR1111 | *unc-13(e51) I; pha-1(e2123) III; him-5(e1490) V; syEx1249[Psrh-142::GCaMP3; pha-1(+)]* |
| UR1112 | *unc-13(e51) I; him-5(e1490) V* |
| UR1113 | *unc-13(e51) I; daf-22(m130) II; him-5(e1490) V* |
| UR1114 | *srd-1(eh1) II; him-5(e1490) V* |
| UR1115 | *tph-1(mg280) II; him-5(e1490) V* |
| UR1131 | *him-5(e1490) V; fsEx527[1.5kb MAB-3::GFP (pDZ162) + cc::GFP line 2]* |
| UR1185 | *mab-3(e1240) II; him-5(e1490) V; fsEx547[srh-142p::mab-3::mCherry; Pelt-2::gfp line 1]* |
| UR1186 | *mab-3(e1240) II; him-5(e1490) V; fsEx548[srh-142p::mab-3::mCherry; Pelt-2::gfp line 2]* |
| UR1187 | *mab-3(e1240) II; him-5(e1490) V; fsEx549[srh-142p::mab-3::mCherry; Pelt-2::gfp line 4]* |
| UR1188 | *mab-3(e1240) II; him-5(e1490) V; udEx212[Pelt-2::gfp; Psrh-142::gfp; Psrh-142::ced-3(p15); Psrh-142::ced-3(p17) line 1]* |
| UR1189 | *mab-3(e1240) II; him-5(e1490) V; udEx213[Pelt-2::gfp; Psrh-142::gfp; Psrh-142::ced-3(p15); Psrh-142::ced-3(p17) line 2]* |
| UR1190 | *mab-3(e1240) II; him-5(e1490) V; fsEx479[Psrh-142::fem-3::mCherry; Pelt-2::gfp line 2]* |
| UR1195 | *him-5(e1490) V; fsEx527[1.5kb MAB-3::GFP (pDZ162) + cc::GFP line 2]; fsEx478[Psrh-142::fem-3(+):SL2::mCherry; Pelt-2::GFP line 1]* |

**Table S2.**
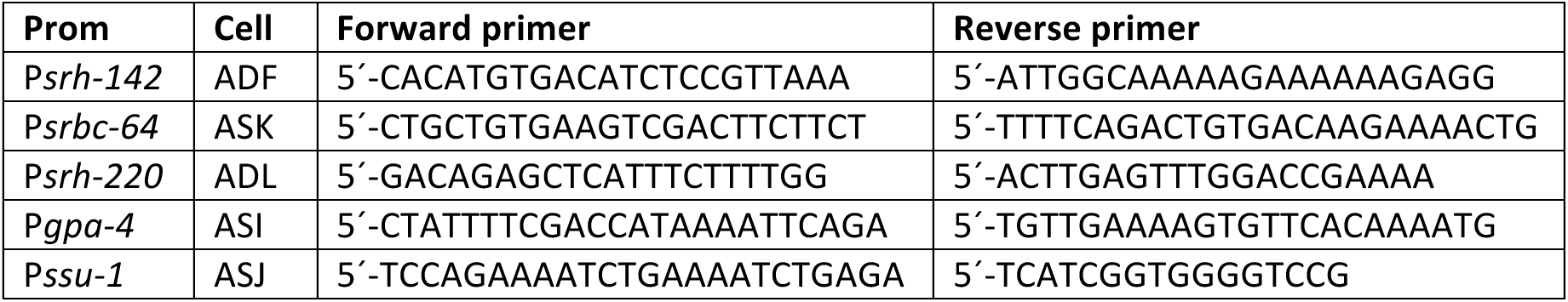
Primers used.

**Figure S1.**
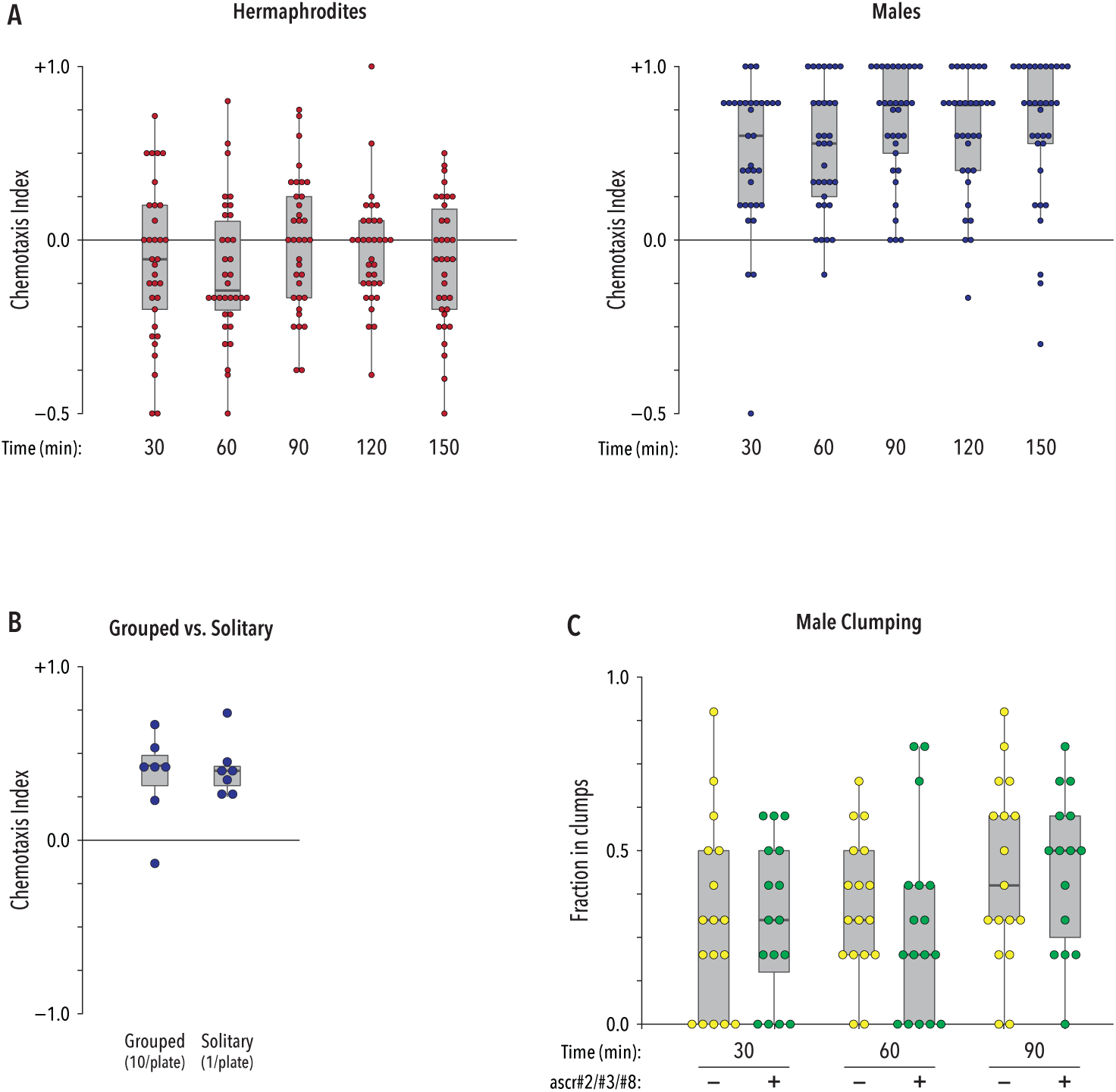
(A) Chemotaxis indices for wild-type animals at 30-min intervals. Over the course of 150 minutes, there was no evidence of adaptation to the ascaroside stimulus. (B) Males tested singly in the quadrant assay behaved similarly to those tested in groups of ten (in the former case, 10 plates were used to calculate each CI value). (C) The presence of ascarosides had no detectable effect on the propensity of males to form clumps.

**S2.**
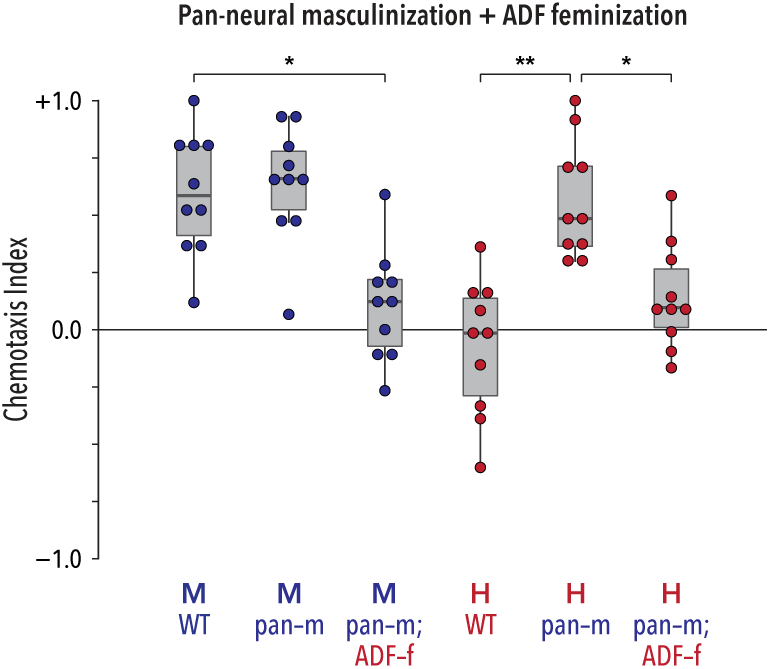
The attraction to ascarosides generated by pan-neural masculinization in hermaphrodites (pan-m) was suppressed by simultaneous feminization of ADF (ADF-f). The effects of feminization are expected to dominate in this case, because expression of the feminizing transgene *tra-2(ic*) in ADF should inhibit the effects of the masculinizing *fem-3(+)* transgene. In males, pan-neural masculinization had no apparent effect on behavior, but as expected, ADF feminization eliminated pheromone attraction.

**Figure S3.**
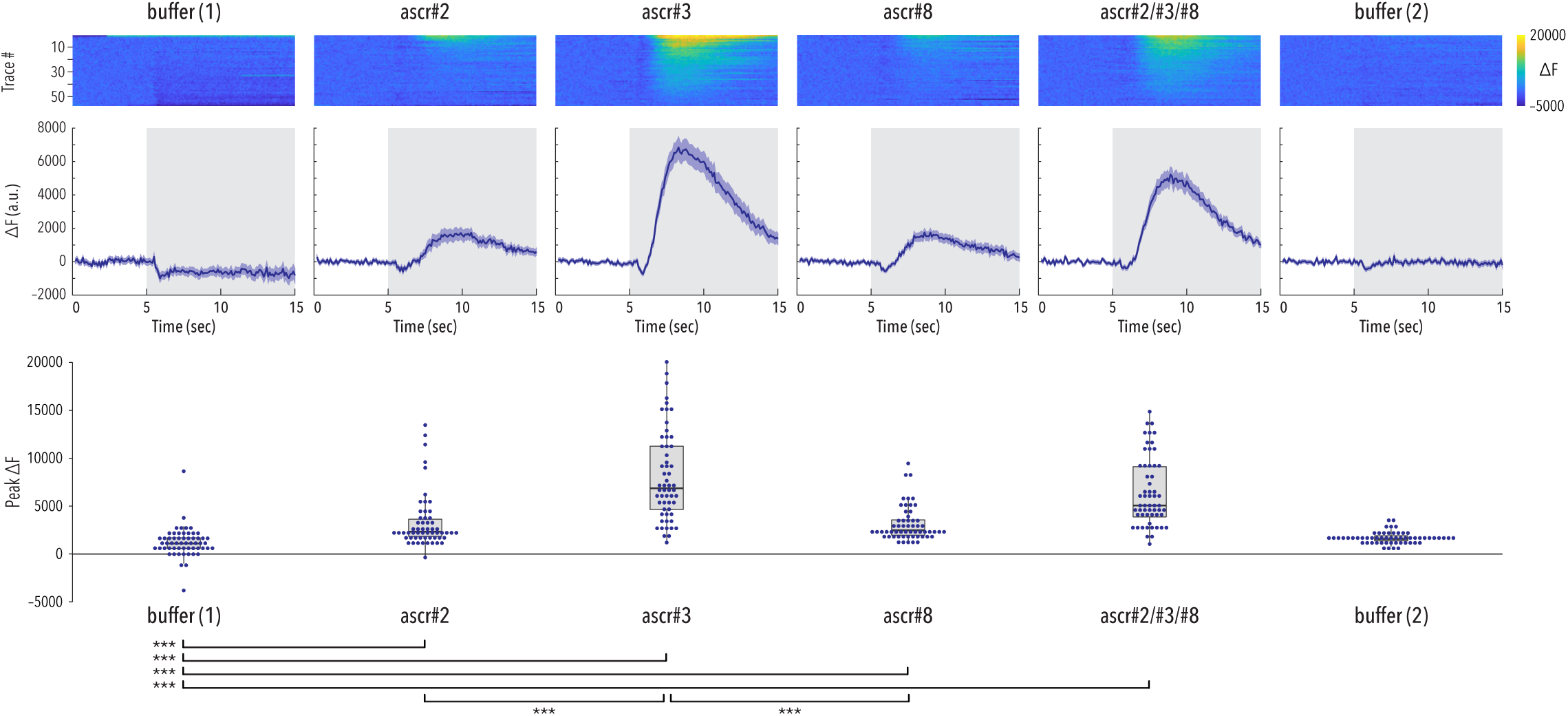
Recordings of ADF calcium transients (as changes in GCaMP3 fluorescence, ΔF) in wild-type males in response to two sequential pulses of buffer, individual ascarosides, an ascr#2/#3/#8 mixture, and buffer. *Top panels* depict pseudocolored graphs of individual neuronal responses. Each row shows a single response; traces are sorted by peak activity. The heatmap represents ΔF. *Center panels* show average ΔF intensities for the recordings depicted in the upper panels. Trace shading indicates +/− SEM. Gray shading indicates the pulse of ascaroside stimulation. *Bottom panels* show dot-plots of peak ΔF values for each recording. Stimulation with ascr#2 or ascr#8 alone elicited a weak calcium response, while stimulation with ascr#3 triggered a markedly stronger response. Because ascaroside stimuli were delivered successively to the same population of animals in the stimulus chamber, some adaptation may have occurred during the stimulation period. This limits the ability of strong quantitative conclusions to be drawn across compounds as compared to naive animals; however, the robust responses to the ascr#2/#3/#8 mixture strongly suggest that any adaptation between stimuli that may have taken place is relatively minor. *n*=57 trials (19 animals, three pulses per stimulus per animal).

**Figure S4.**
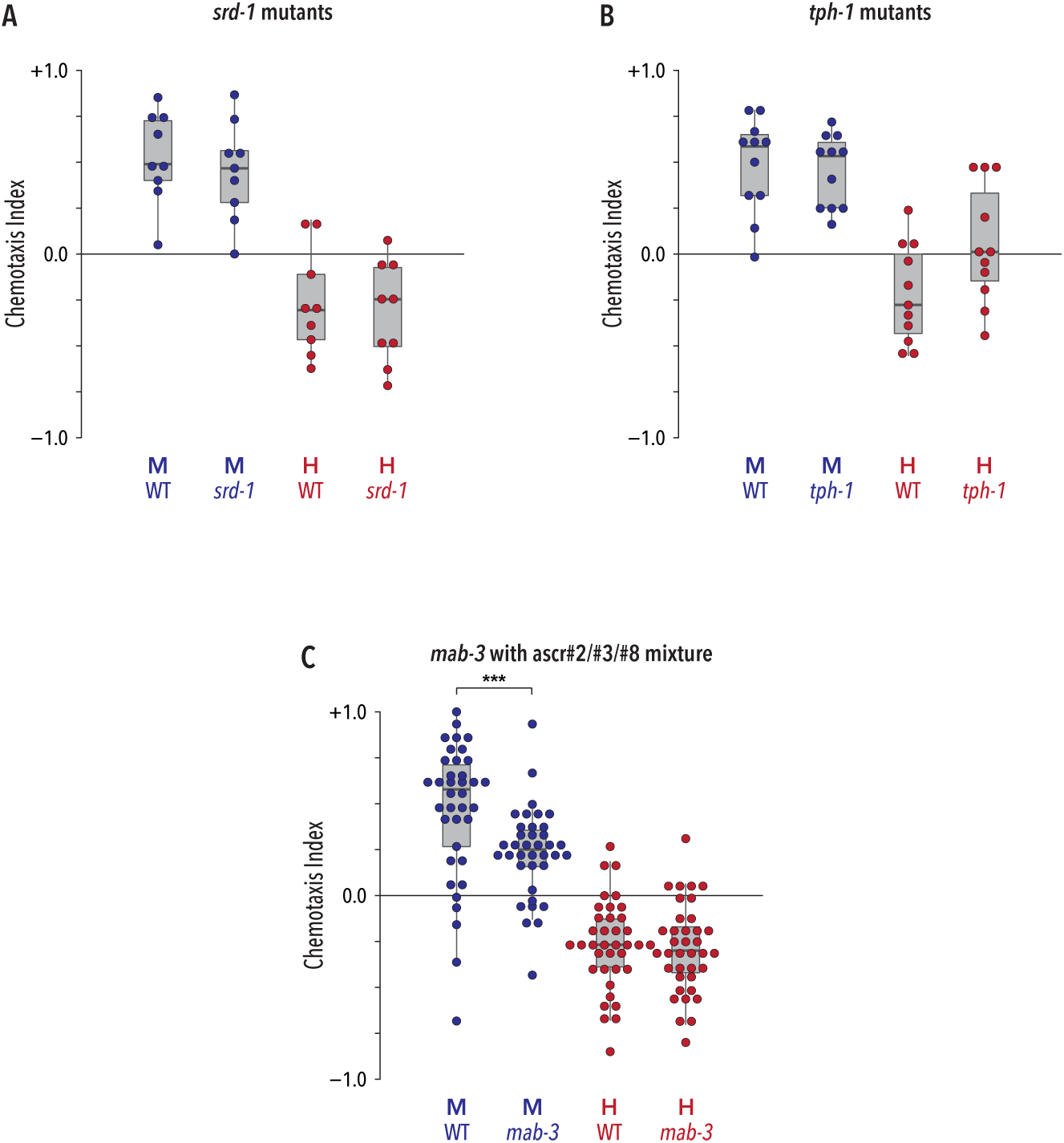
(A) The behavior of *srd-1* mutants in the quadrant assay was indistinguishable from wild-type. (B) Loss of the serotonin-biosynthetic enzyme *tph-1* had no detectable effect on the behavior of males or hermaphrodites. (C) *mab-3* mutant males exhibited a partial loss of attraction to the ascr#2/#3/#8 blend, comparable to the effects of *mab-3* loss on attraction to ascr#3 presented individually (see Fig. 4a). *mab-3* mutant hermaphrodites behaved similarly to wild-type hermaphrodites.

**Figure S5.**
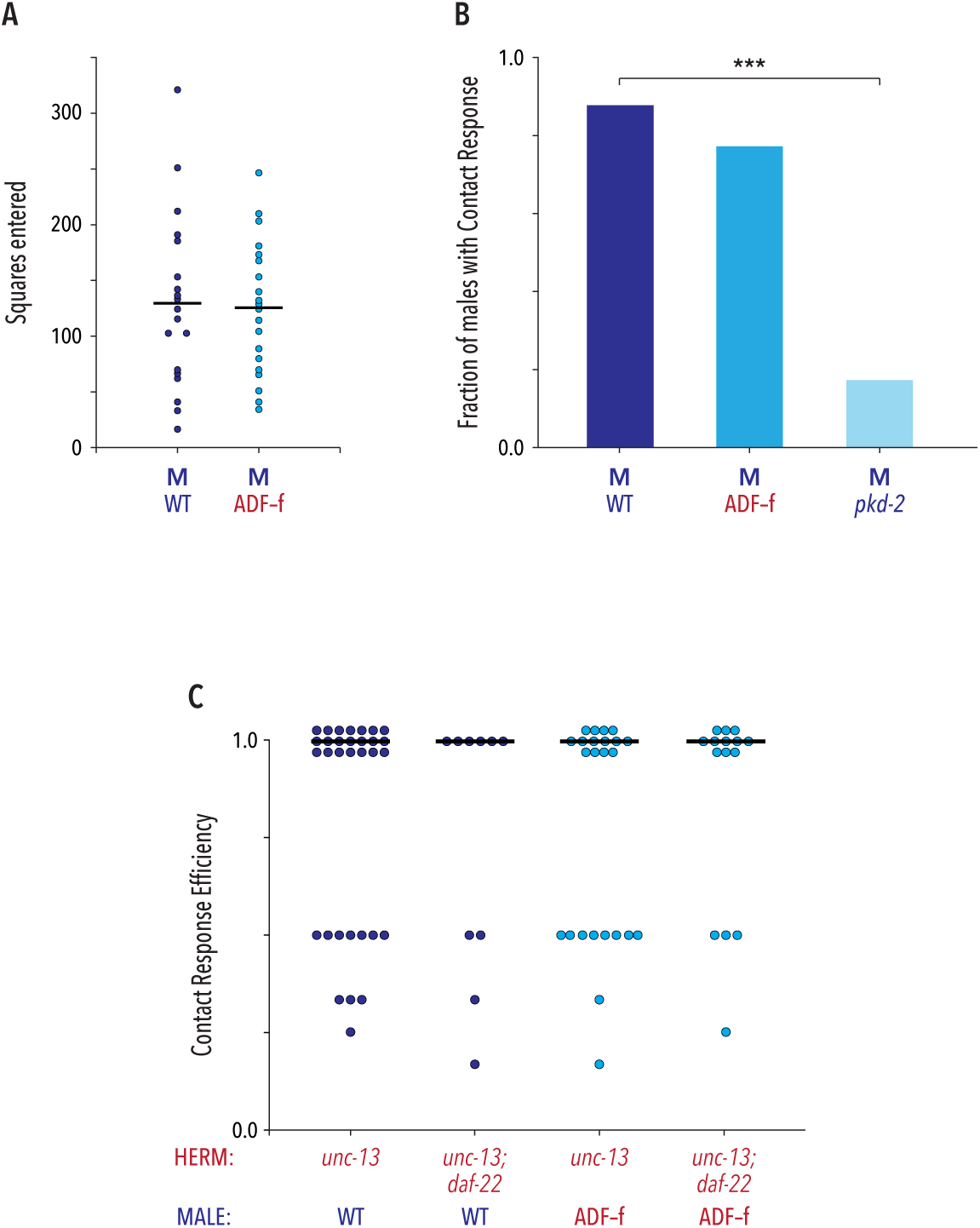
(A) Male exploratory behavior, measured as the fraction of squares in a grid entered over 60 minutes, was not noticeably affected by ADF feminization (ADF-f). (B) ADF feminization did not detectibly disrupt the frequency of contact response, the first step of male mating behavior. *pkd-2* mutant males, whose response behavior is severely compromised[109], were used as controls. (C) Contact response efficiency, scored as the reciprocal of the number of physical encounters until response occurred (see Methods), was not detectibly affected by ADF feminization, nor did it depend on the ability of hermaphrodites to produce ascarosides (*unc-13* vs. *unc-13; daf-22*).

## ACKNOWLEDGEMENTS

We are grateful to Denise Ferkey and Michelle Krzyzanowski for *Pelt-2::gfp* and the ADF ablation strains; to Alon Zaslaver for the *ADF::GCaMP3* transgene; and to Jan Kubanek for help with the Parallel Worm Tracker script. We thank Jagan Srinivasan for sharing unpublished data and for helpful feedback, and Ilya Ruvinsky for sharing unpublished findings. We are grateful to Clare McMahon and Kelly Rineer for contributing data to Figs. S3 and 5, respectively. We thank members of the Portman lab and the Western New York Worm Group for thoughtful discussion. Some strains were provided by the *Caenorhabditis* Genetics Center, which is funded by NIH Office of Research Infrastructure Programs (P40 OD010440). This work was funded by NIH NRSA F31 NS086283 (K.F.), NSF IOS 1353075 (D.P.), NIH R01 GM108885 (D.P.), NSF CBET 1605679 (D.A.), NSF EF 1724026 (D.A.), and Burroughs Wellcome Fund CASI Award (D.A.).

## AUTHOR CONTRIBUTIONS

Conceptualization, K.A.F., J.L., R.C.L., D.R.A., and D.S.P.; Methodology, K.A.F., R.C.L., D.R.A., and D.S.P.; Investigation, K.A.F., J.L., and R.C.L.; Resources, F.C.S.; Writing — Original Draft, K.A.F., R.C.L., and D.S.P.; Writing — Review & Editing, K.A.F., R.C.L., J.L, F.C.S., D.R.A., and D.S.P.; Supervision, D.R.A. and D.S.P.; Funding Acquisition, K.A.F., D.R.A., and D.S.P.

